# Cell size and shape regulation of *E. coli* determines surface area scaling with volume

**DOI:** 10.1101/2022.03.23.485554

**Authors:** Tanvi Kale, Dhruv Khatri, Chaitanya A. Athale

## Abstract

The scaling of surface area and volume of cells has widespread consequences for cell physiology, growth and adaptation. While the surface area increases with volume as SA ~ V^γ^ the scaling exponent for proportional growth maintaining the shape and aspect is γ ~ 2/3 or 0.66. However most well-studied cellular systems deviate from this standard exponent. At the same time, a mechanism that could predict the biological or physical basis of these scaling relations remains unclear. Here, we quantify the surface area scaling with volume of *Escherichia coli* cells with varying growth rates and under different conditions and find the scaling exponent varies from γ ~ 0.7 to 0.9. A model of uncorrelated statistical variation of cell lengths and widths can reproduce the exponent observed in experiment. Average values of length and width on the other hand results in an impression of ‘ideal’ geometric scaling, as reported in some studies. Experimental data however suggests that *E. coli* cell width is strongly correlated with length and a model of saturation best explains the observations. We hypothesize this model of cell size and shape regulation could serve the function of optimizing flux of nutrients, within the constraints of the cell division machinery.

## Introduction

Cell shape and size are recognizable characteristics of cellular life forms. Bacterial cell shape and size are used classically to identify them. The regulation of bacterial cell shape has in the past been considered to influence surface area and volume, with rod-shaped cells having an advantage over simple spherical cells (Koch, 1995). At the same time why a certain cell attains a size is considered an outcome of evolution constrained by the molecular machinery driving it (Bonner, 2006). Cell sizes of bacteria also change with growth rate as summarized by the Schaechter’s nutrient growth law (Schaechter et al., 1958). The mechanism for this has been more recently been identified as OpgH in *Escherichia coli* (Hill et al., 2013) and its functional homolog UgtP found in Bacillus (Weart et al., 2007). At the same time, even in a population of synchronized cells, cell size is seen to follow a distribution, attributed to either model predictions of division and growth variability (Hosoda et al., 2011) or single cell cycle time variation (Osella et al., 2014). In previous work, we found *E. coli* cell length variability to also increase with growth rate, that could be explained by a combination of stochasticity in DNA replication and inhibition of cell division resulting in increased cell filamentation (Gangan and Athale, 2017). While multiple mechanisms can result in *E. coli* cell filamentation in a population, whether such heterogeneous cell size distributions could have a functional role for populations remains unclear.

The stereotypical size and shape of single *E. coli* cells has driven the discovery of multiple homeostatic mechanisms that combine the FtsZ-ring assembly and divisome regulation (Goehring and Beckwith, 2005; Wang et al., 2005), the MinCDE system that regulates the position of the division site (Varma et al., 2008) and a parallel nucleoid occlusion pathway that prevents division until the nucleoid segregates (Bernhardt and De Boer, 2005; Cambridge et al., 2014; William Adams et al., 2015). Of the many phenomenological models of bacterial cell size regulation, a model of a constant increment or ‘adder’ at every cell cycle (Amir, 2014) is best supported by the evidence of correlation between size at birth and the addition of size at division (Campos et al., 2014; Taheri-Araghi et al., 2015). Recent analysis has shown that the increment in volume best explains the data, while measurement errors in cell length and cell size variability could result in alternative models being equally likely (Facchetti et al., 2019). While most models focus on the average size and shape, variability is important in developing models.

The growth of surface area with volume for a geometric solid is expected to scale with a scaling exponent, γ of 2/3 or 0.66. Cell shape however does not appear to follow such ideal scaling with a higher exponent observed, explained in part by the diffusion limited nature of nutrient uptake (Koch et al., 1996). Nutrient uptake in turn is an important selective feature driving specification of cell shape in bacteria (Young, 2006). Indeed the robustness of *E. coli* cell shape is demonstrated by the reversion of *E. coli* to typical aspect ratios of length:width in a 50,000 generation long term evolution experiment (LTEE) experiment (Grant et al., 2021). This optimality could be the result of the constraints governing the rate of surface area addition relative to volume growth rate that sets cell size and aspect ratio (Harris and Theriot, 2016). A recent set of studies proposed gram-negative *E. coli* surface area follows ideal geometric object scaling with volume, 2/3, while *Bacillus subtilis* deviates from it (Ojkic et al., 2019), and a growth rate model coupled to FtsZ assembly fit to experimental data was developed to explain this scaling (Ojkic and Banerjee, 2021). This suggests contradictions in the literature reporting on the quantitative effect of growth rate on cell size and shape of *E. coli*.

In this study, we proceeded to quantify the cell lengths and widths of populations of *E. coli* populations. We find the analysis of surface area scaling with volume from membrane stained cells, growth dependent changes in cell size and mutants for nucleoid occlusion and replication stalling coupled to division, are all greater than the geometric exponent 2/3 (or 0.66), ranging instead between 0.7 and 0.9. We show that variability of cell lengths and widths can partially explain this deviation based on statistical sampling. We examine three alternative models of coupling between cell widths and lengths, and find only one model of increase and saturation of width can explain the data. We attempt to reconcile this with the scaling of SA with V and use it to predict the functional role such scaling could serve.

## Results

### Surface area scaling with volume of rod-shaped bacterial cells and their size variability

The size and shape of a cell determines the nature of allometric scaling. Surface area (SA) is expected to scale with volume (V) based on the expression:

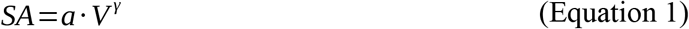

where γ is the scaling exponent. For simple geometric shapes with proportionate increase in dimensions, γ ~ 2/3 (0.66). This is also true of rod-shaped cells if the condition is met that the aspect ratio (AR), i.e. the ratio of length to width, is constant (**Figure 1A**). However, if the width of such a cell is maintained constant, driven by mechanics of the either intracellular cytoskeleton or cell wall, then the scaling exponent γ will change to 0.96. In previous work comparing cell shape scaling differences between *Bacillus subtilis* and *Escherichai coli* had reported that *E. coli* would scale with an exponent of 0.6 while *B. subtilis* would scale with an exponent of 0.9. However, for such a model to be validated, *E. coli* would have to maintain a constant aspect ratio. At the same time, a population of *E. coli* displays natural cell size variability that is the cumulative result of multiple mechanisms at a single cell level including gene expression stochasticity (Elowitz et al., 2002), variability of molecular partitioning at division (Rosenfeld et al., 2005), replication stochasticity and coupling to division (Gangan and Athae, 2017) and DNA damage response (Raghunathan et al., 2020). As a result, we hypothesize that these mechanisms result in less dramatic changes in width as *E. coli* cell length increases due to either stochasticity in replication, division or segregation of proteins (**Figure 1B**). Combined with previous observation of a wide distribution of cell lengths, we therefore expect that the scaling exponent of *E. coli* populations is likely to deviate from that of simple geometric objects.

**Figure 1.**
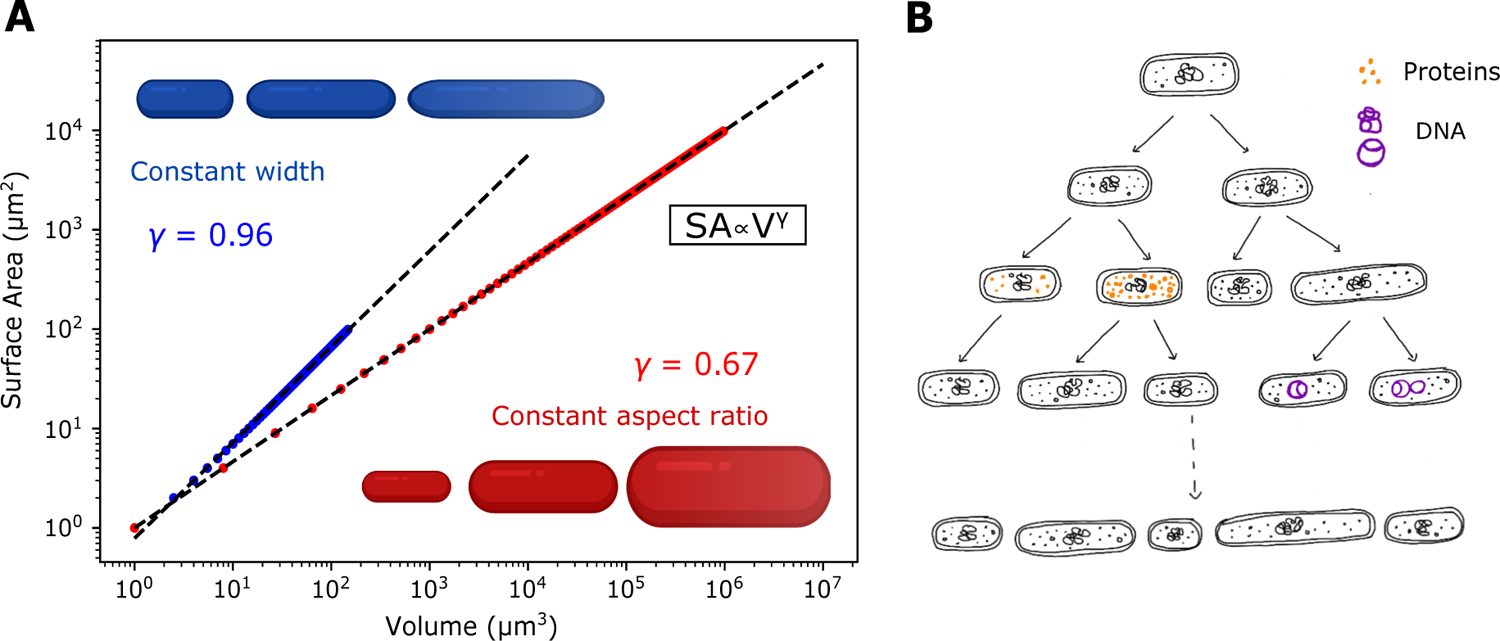
Cell size results in differences in allometric scaling of rod-shaped bacteria. **(A)** Rod shaped bacteria were simulated with a range of lengths from 1-100 μm and widths either based on w=L/A_R_ (red) or w=c (blue) resulting in increasing surface area (SA)with volume (V). The data was fit to SA ~ V^γ^ with γ as the fit parameter, as noted in the graph. Here, A_R_ = 5.16 and c = 1 μm. **(B)** Schematic of a multiple rounds of bacterial cell division highlighting asymmetric protein segregation (orange) or DNA replication stochasticity (purple) as possible mechanisms that result in population cell length variability.

### SA-V scaling of membrane stained *E. coli* cells deviates from the 2/3 law

In order to test the hypothesis of scaling of SA with V for cells of varied lengths, we proceeded to measure *E. coli* cell length and width distributions. Mid-log phase cells of *E. coli* MG1655 untreated or treated with 100 μg/ml cephalexin were compared to DH5α cells based on cell membrane stained with FM4-64 (**Figure 2A,B**). An image analysis pipeline that detects the cell boundaries and uses median cell width statistics resulted in cell length and width measures (**Figure 2C**). Consistent with previous reports, *E. coli* MG1655 cells have narrower spread of distribution compared to DH5α, while cephalexin treatment increases mean lengths and their spread compared to untreated (**Fig. 2D**). The width on the other hand of both strains appear to be similar, resembling a normal distribution with the mean width of cephalexin treated cells slightly higher than the others (**Figure 2E**).

**Figure 2.**
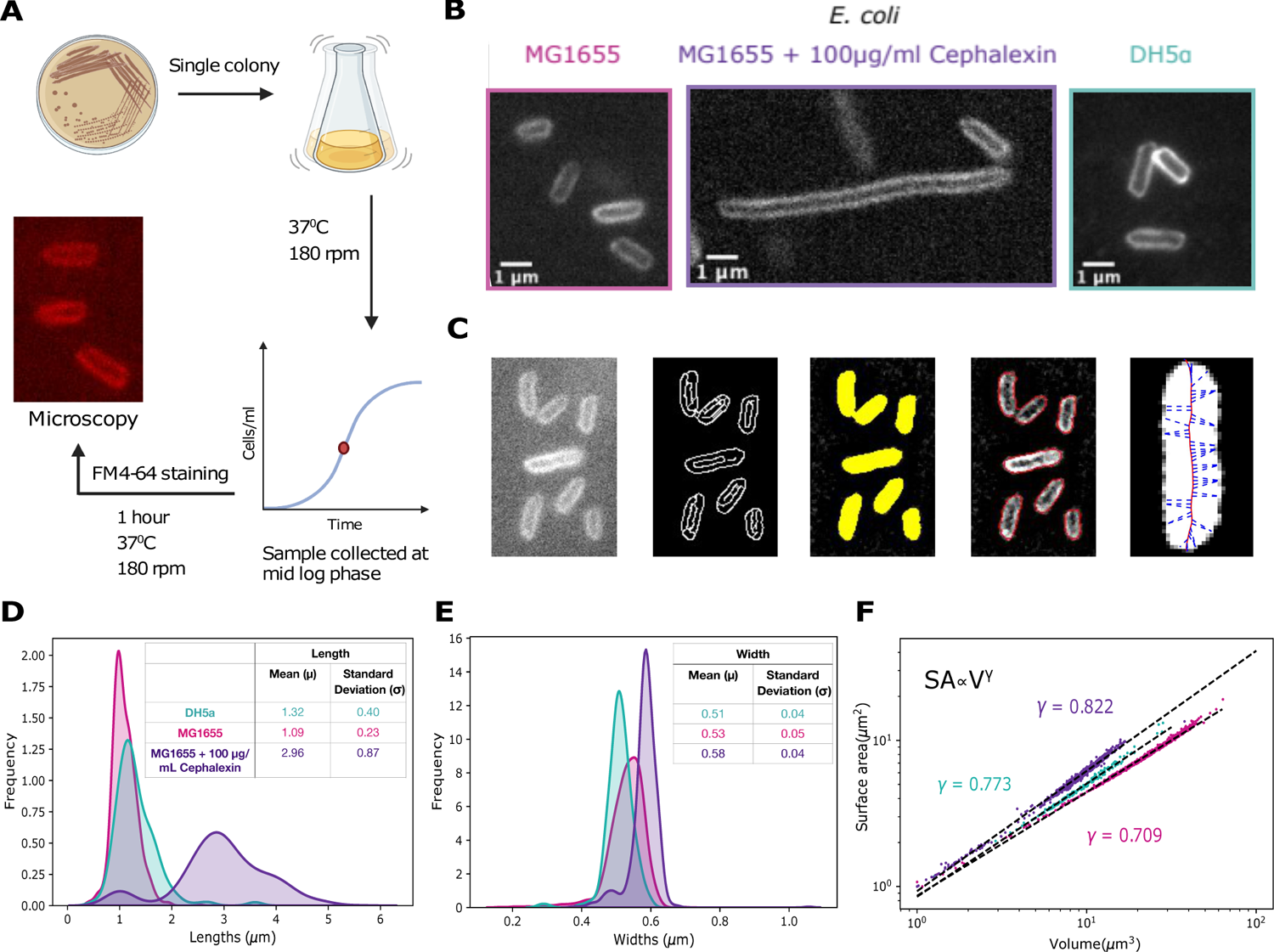
Quantification of *E. coli* cell lengths and widths and SA-V scaling. **(A)** The schematic represents the method used to stain *E. coli* cells with FM4-64 sampled from mid-log phase cultures. **(B)** *E. coli* MG1655 either untreated (magenta) or treated with 100 μg/ml cephalexin (purple) and DH5α (cyan) were imaged in fluorescence microscopy. Box colors: strain or treatment. **(C)** Cell lengths and widths were measured using a morphometric image analysis pipeline as described in the Materials and Methods section. **(D)** The resulting cell length and **(E)** width distributions are plotted. **(D, E** Inset) Tables represent the respective distribution mean and standard deviations. (**F)** The surface area (SA) and volume (V) calculated from the lengths and widths (see Materials and Methods) were normalized by the minimal surface area (SA_0_) and volume (V_0_) and the data fit to SA = a*V^γ^ resulting in γ = 0. 83. Coloured circles indicate- magenta: *E. coli* MG1655, purple: *E. coli* MG1655 + Cephalexin, cyan: *E. coli* DH5α. The schematic was created with BioRendnder.com.

The surface area, SA and volume of the cells calculated based on geometric expressions (Equations 6,7, Materials and Methods) were normalized by the respective minimal values (SA_0_, V_0_) and fit to the scaling relation in Equation 1 **(Figure 2F)**. The SA-V of the three different conditions appears to scale with an exponent γ of 0.83, deviating from the ideal geometric value of 0.66. In order to examine if this is consistent across other conditions of lab strains, we proceeded to re-examine the scaling of cell populations with varied growth rates.

### Effect of growth rate and nucleoid occlusion mutations on scaling of *E. coli* SA-V scaling in populations

The observed increase in mean bacterial cell size with growth rate was formulated as Schaechter’s nutrient growth law (Schaechter et al., 1958) and more recent single-cell measurements of *E. coli* mean cell length and width have confirmed it (Taheri-Araghi et al., 2015). However, growth rate results in not just changing *E. coli* averages of cell size, but also their population variability as we have reported previously (Gangan and Athale, 2017). Cell sizes were taken from mid-log phase *E. coli* cells whose growth rate had been modulated by the nature of the medium and carbon source (**Figure 3A,B**), with mean length and width as a function of growth rate (*r*) were fit to a previously described model (Taheri-Araghi et al., 2015) where:

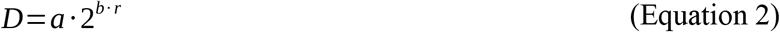

where the dimension *D* corresponds to either length (*L*) or width (*w*). The fit parameters of length (a = 1.73 and b = 0.39) are consistent with microfluidics data. We therefore use fit obtained for width by Taheri-Araghi et al. (2015) to model width as w=0.41*2^0.33*r^ generating predicted values for each growth rate (**Figure 3C**), to arrive at SA and V values. Surface area of growth rate modulated data then scales with volume with an exponent γ of 0.88 for all conditions (**Figure 3D**). Individual growth rate SA-V data fit by the scaling relation (**Figure S1**) demonstrates a small but systematic increase in γ with growth rate, while the x-intercept of the plot, *a* ~ 2π as expected from geometry (**Figure 3E**). Analyzing the SA-V scaling of *E. coli* strains mutated for the replication processivity factor, recA, its effector sulA and nucleoid occlusion factor, slmA (Gangan and Athale, 2017) reveals an exponent ranging between 0.8 and 0. 9 (**Figure S2**), consistent with their role in cell elongation compared to wild type (**Figure 3B**).

**Figure 3.**
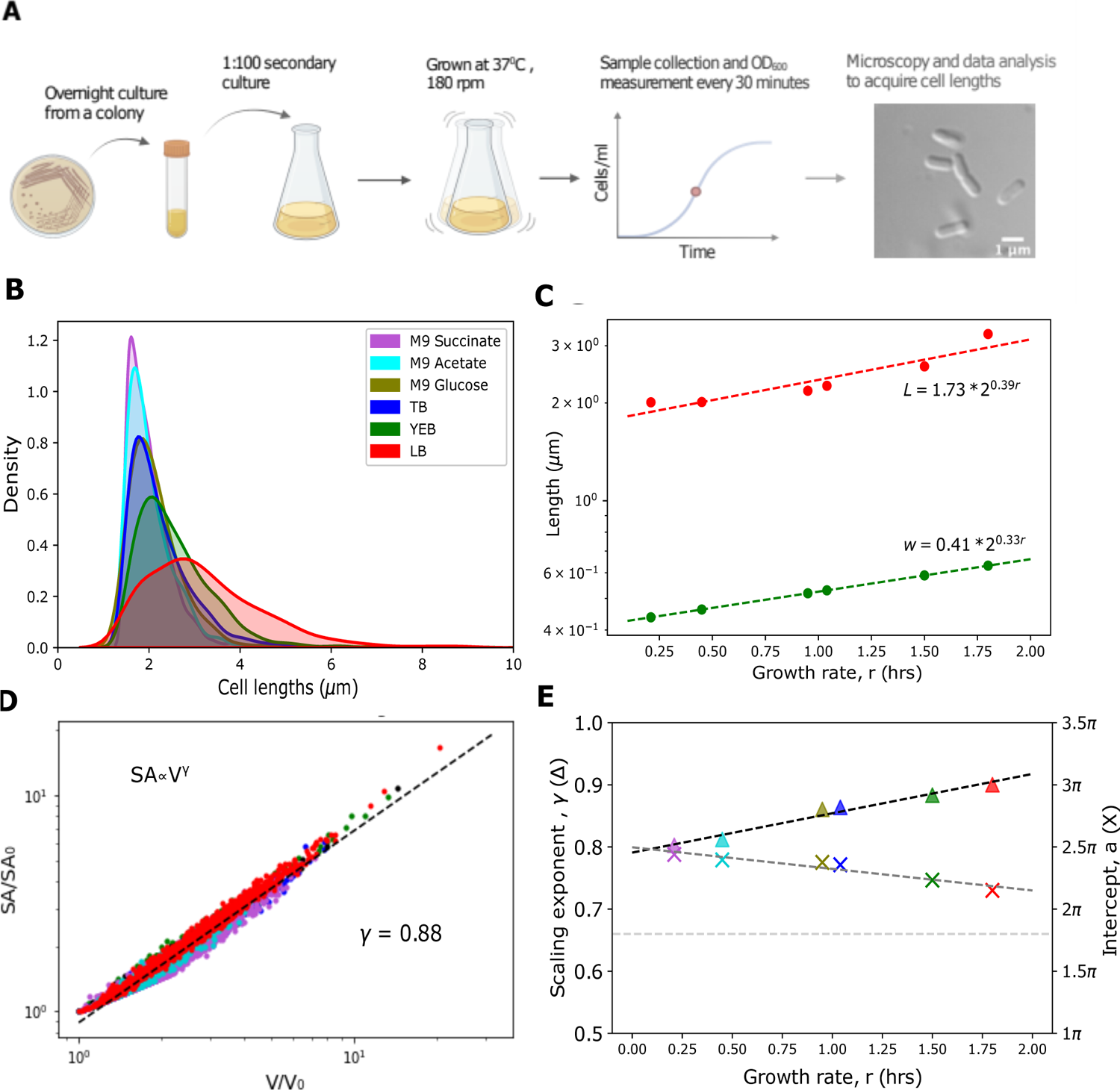
Growth rate dependent surface area scaling with volume of *E. coli* from DIC images. **(A)** *E. coli* MG1655 cells were sampled from mid-log phase cultures and imaged in DIC, as shown schematically. **(B)** Cell length distributions from cultures in different growth media (legend) are plotted based on previously described work (Taheri-Araghi et al., 2015). **(C)** The mean cell length (L, red circles) and width (w, green circles) are plotted as a function of the growth rate (r), which was modulated by changing the growth medium. The cell lengths (red circles) were fit to Equation 2 (see Results section) with fit parameters a = 1.73 and b = 0.39, while widths were calculated from Equation 1 solved for width with a= 0.41 and b= 0.36, based on previous work (Gangan and Athale, 2017). Dashed line: model. **(D)** The SA and V values normalized by their corresponding minimal values (SA_0_, V_0_) were fit to Equation 1 resuling in γ = 0.88. Colors indicate growth media. **(E)** The growth rate dependence of the (*left y-axis*) scaling exponent, γ (▲) and (*right y-axis*) the y-intercept of the fit (x), i.e. the value of the normalized surface area area SA/SA_0_ when V/V_0_ = 1 in units of π are plotted. Dashed lines indicate linear fits to the data. Colors- red: LB, green: YEB, blue: TB, olive-green: M9+ Glucose, cyan: M9+ Acetate and purple: M9+ Succinate. The schematic was created with BioRender.com.

This demonstrates that *E. coli* SA-V scaling exponent exceeds the 2/3 value claimed in recent reports in literature (Ojkic et al., 2019; Ojkic and Banerjee, 2021) independent of growth conditions, mutant background or method of sample preparation. It however remains unclear how the value reported earlier could emerge- whether it is simply a consequence of having ignored statistical variability or a physiological mechanism that regulates bacterial cell size regulation.

### Statistical averaging can explain the emergence of 2/3 scaling of SA-V

Statistical variability is apparent in most populations of *E. coli* grown even under strictly synchronized conditions. We hypothesize that cell lengths and widths are subject to uncorrelated randomness in the underlying molecular processes. We test this hypothesis by two approaches - experimental data and statistical simulations. As a first step, the volume and surface area distributions for each growth rate were averaged and the SA with V fit to the scaling relation. To our surprise the scaling exponent from population data of 0.88 changes to 0.69 (**Figure 4A**), closer to the ‘ideal’ scaling exponent of 0.66. This appears to demonstrate that distributions, as opposed to averages, strongly influence the scaling relation and inferences made based on them. In order to address the generality of this result, as a second step, we simulated rod-shaped cells whose cell lengths and widths varied while maintaining a constant aspect ratio, i.e. proportional length and width. Cell lengths were sampled from lognormal distributions with increasing means from 1.1 to μm and standard deviations of 0.22 to 2.53 μm respectively (**Figure 4B**), while widths were sampled from a normal distribution with the mean width inferred from the ratio L/A_R_ with aspect ratio A_R_ = 5.16 (**Figure 4C**), and randomly combined pairwise to calculate SA and V. The aspect ratio was taken from average values reported previously (for details refer to Materials and Methods). The surface area data with volume scaled with exponents γ ranging from 0.65 to 0.93, for increasing average cell sizes (**Figure 4D**). The increasing mean lengths and widths, represent the effect of growth rate. The choice of distribution did not affect the outcome since similar results were obtained when lengths were sampled from a normal distribution (**Figure S3**).

**Figure 4.**
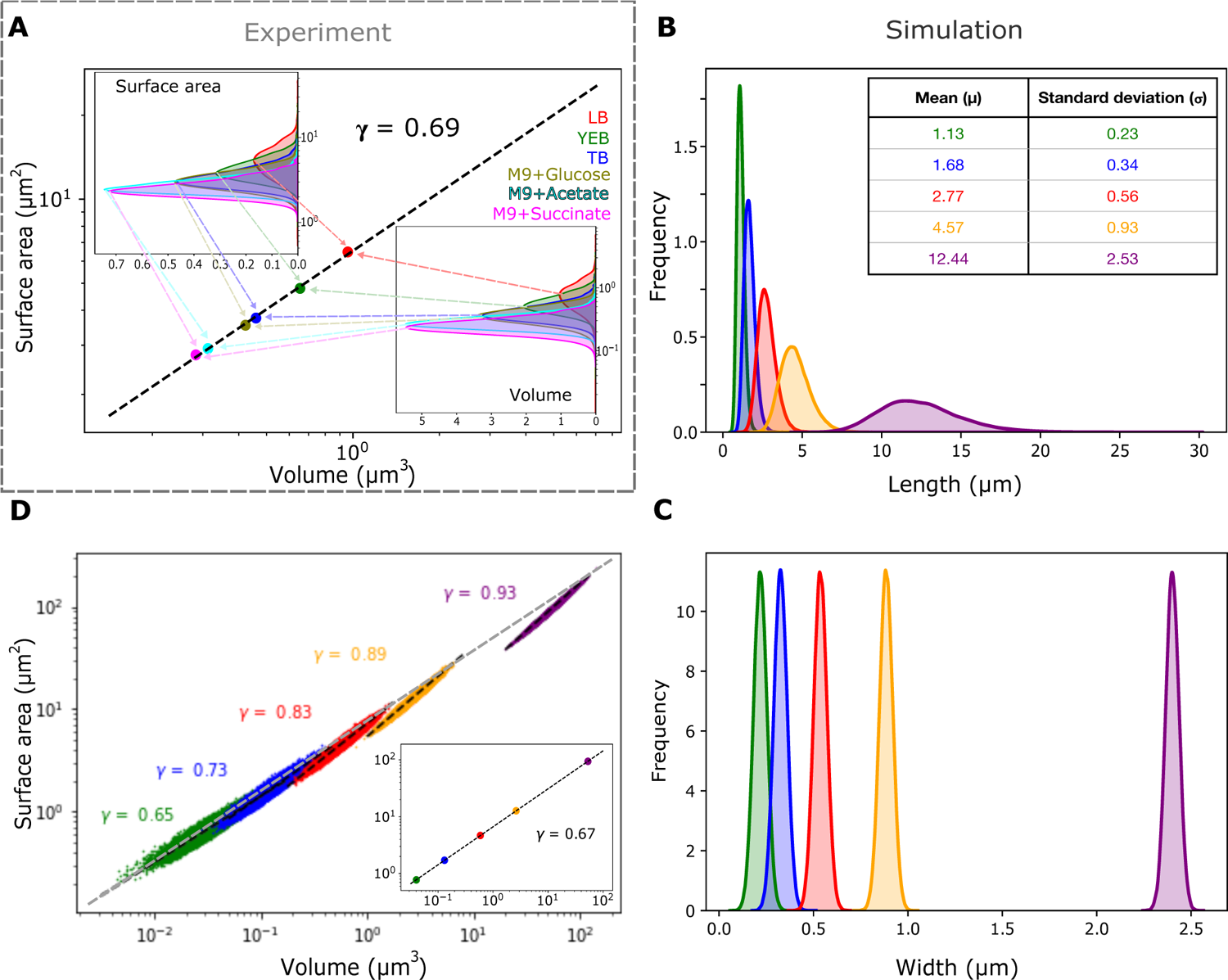
The effect of cell size population variability and averaging ons allometric scaling. **(A)** The surface area and volume averages (circles) of *E. coli* MG1655 taken from their respective distributions were fit to Equation 1 (Results section) to infer the scaling exponent γ. *Inset:* Frequency distributions of surface areas (top) and volumes (bottom). Colors- Growth media. Red: LB, green: YEB, blue: TB, olive: M9+ Glucose, cyan: M9 + Acetate and purple: M9+ Succinate. **(B, C)** Simulated distributions of lengths and their corresponding widths (colors) with random sampling from **(B)** lognormal distributions of lengths with arithmetic mean (μ_L_) and s.d. (σ) (inset: mean and s.d.) and **(C)** normal distributions of cell widths. Width distributions are based on mean width μ_w_= μ_L_/A_R_ and a constant standard deviation (from Figure 2E and Materials and Methods), while the aspect ratio A_R_ = 5.16. **(D)**The resultant surface area and volume for each of these length and width distributions (coloured circles) were fit to Equation 1 to estimate γ for each dataset (colors). *Inset:* The mean surface area and volume for the same datasets fit to Equation 1 resulting in γ = 0.67.

Thus we find that population statistics of cell size are critical for the estimation of SA-V scaling exponents. However, it remains unclear if uncorrelated sampling of cell lengths and widths is a valid assumption. Therefore we proceed to examine the relationship between lengths and widths, which could reveal potential regulation of *E. coli* cell size.

### Models of coupled increases in cell length and width of E. coli

The role of a cell-size regulation in scaling requires us to address how exactly the length and width of *E. coli* cells may be coupled. To address this, we assume three alternative non-molecular (phenomenological) models of *E. coli* size regulation (**Figure 5A**). The simplest model, **model I**, assumes the cell width (*w*) is held constant due to potential molecular interactions of the cell wall and cytoskeleton resulting in:

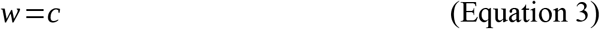

where c is a constant determined by some intrinsic limits to cell width. Such a limit could be potentially set by the interplay of membrane precursors and cytoskeletal elements like MreB. An alternative **model II**, assumes that the aspect ratio is constant, with length and width linearly related to each other. This model corresponds to *E. coli* growth rate dependent scaling of SA-V with an exponent γ of 2/3 (Ojkic et al., 2019) and a molecular mechanism based on the kinetics of UgtpP monomers and FtsZ assembly (Ojkic and Banerjee, 2021). The resulting linear dependence of width and length, is expressed as:

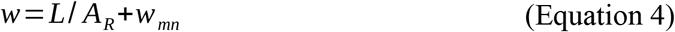

where A_R_ is the constant aspect ratio, L is the cell length and w_mn_ is the minimum width. Such a model would assume cell volume increase is proportionate and resembles ideal geometric objects. However, our observations suggest that there may be an initial rapid increase in width with length but an intrinsic self-limitation does not allow it to exceed a value, that we refer to as the width saturation model, **model III** and it can be written as:

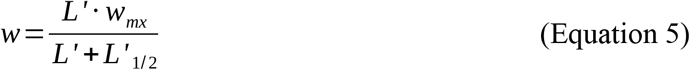

where w_mx_ is the normalized maximal value of the cell width, and L’ and L’_1/2_ are the normalized cell lengths and the length at which the width is half maximal and normalized length is L'=L-L_mn_ where L_mn_ is the smallest length measured. Such a model assumes that cell width increases but has an intrinsic upper limit. Such a limit could be imposed by a molecular mechanism, related for example to the MreB cytoskeleton coupled to the membrane and may act as a sensor and regulator. The *E. coli* cell length variation with width of MG1655 and DH5α strains plotted demonstrates a striking fit to model III (**Figure 5B**). Indeed comparing all three models demonstrates that they predict distinctly different widths as a function of length, for a range of cell lengths (**Figure 5C**). The free parameters of the models such as average cell width (**model I)**, mean aspect ratio of E. coli (**model II**) and the minimum width are taken from a previous studies, while wmx, L1/2 and Lmn (**model IIII**) were estimated from data (**Figures 5B)**. The SA with V values calculated for the three models predict distinctly different SA-V scaling exponents with (I) γ = 0.96 for constant width, (II) γ = 0.67 for the linear model and (III) γ ranging between 0.85 to 0.9 depending on the exact range of lengths (**Figure 5D**). This range of exponents for the saturation model (III) arises due to the fact that for small cells of lengths upto 2 μm the width increases rapidly, but longer cells see no change in width, resulting in a scaling exponent similar to the constant width.

**Figure 5.**
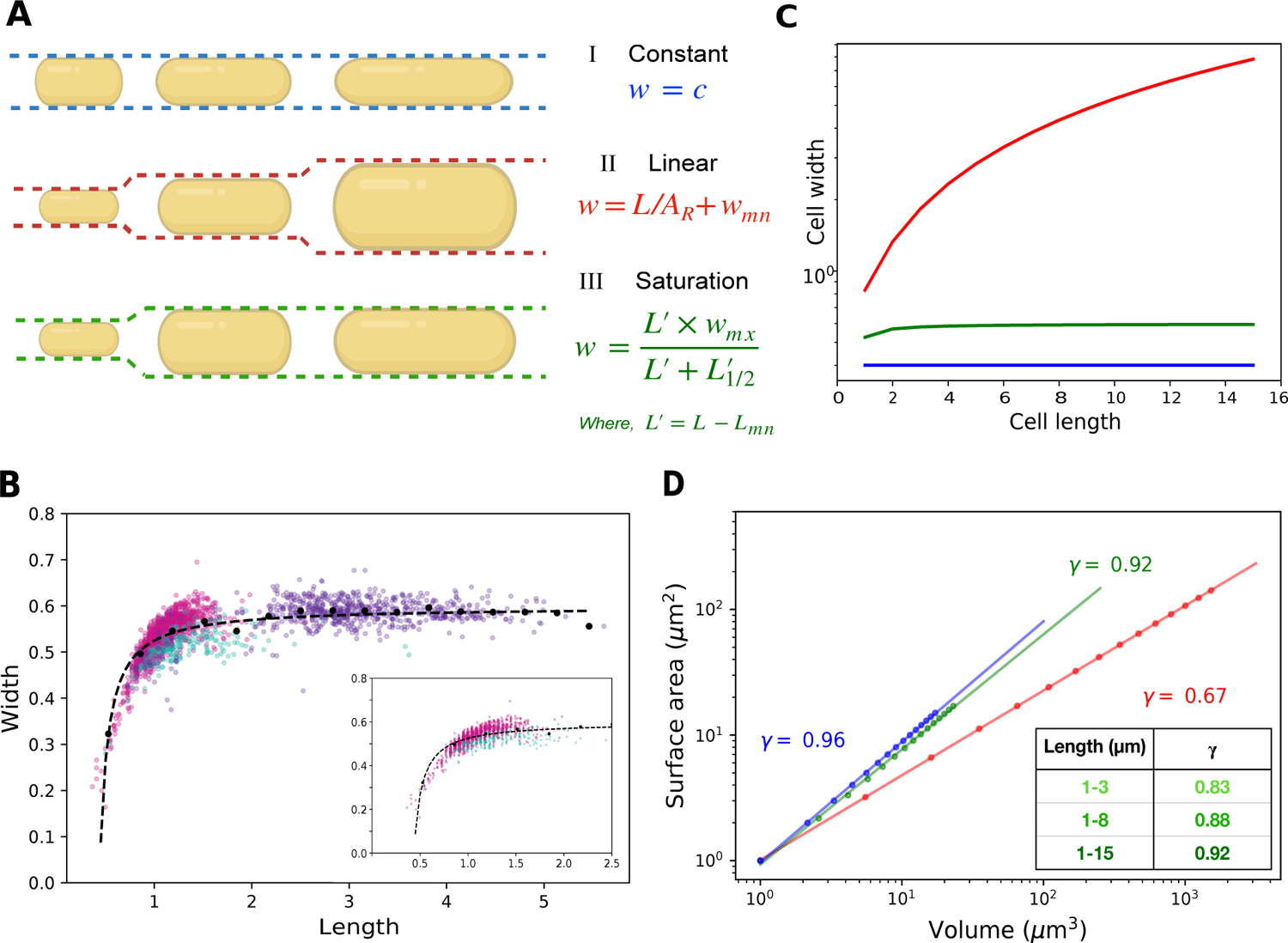
Modeling *E. coli* cell width dependence on length compared to experiment. **(A)** Three phenomenological models of bacterial cell width (w) dependence on cell lengths (L) are depicted with equations and schematics (created with BioRender.com). Model I: constant width (blue), model II: linear dependence on length (red) and model III: saturation (green). **(B)** Experimentally measured *E. coli* cell widths are plotted as a function of cell lengths (colored circles) obtained from microscopy. Binned average data (black circles) were fit to all 3 models. Only model III fit the data well with parameters L_1/2_=0.52 μm, L_mn_=0.44 μm and w_mx_=0.59 μm. Colors indicate: MG1655 untreated: magenta and 100 μg/ml cephalexin treated: purple and DH5α: cyan. (*Inset*) Experimentally measured *E. coli* MG1655-untreated and DH5α cell widths are plotted as a function of cell lengths fit with model III. **(C)** The 3 models predict differing trends of width as a function of length. Model parameters are A_R_=5.16, w_mn_=0.33 μm, constant width = 0.4 μm, L_1/2_ = 0.52 μm. **(D)** The scaling of SA with V (circles) resulting from the three models of width regulation for increasing lengths (I, II, III), were fit to Equation 2 to estimate γ resulting in (**model I**) 0.96, (**model II**) 0.68 and (**model III**): 0.92. (*Inset*) M**odel III** predicts different values of γ for increasing ranges of cell lengths.

Thus we find the proportional model of increasing width with length is falsified based on comparison to experimental data and a width saturation model best cell size data. We proceed to examine if such scaling could have any functional role.

### Constraints on cell width growth due to limitations in surface area synthesis

Previous studies have shown that surface area growth is self limiting as compared to volume growth rate (Harris and Theriot, 2016, 2018; Harris et al., 2014). Since our experimental data of E. coli points to increasing cell lengths but self-limiting widths, we proceeded to examine how this might relate to the surface area for the different models (**Figure 6A,B**). By dimensional analysis we can show that for **model I** (constant width) *SA* ∝ *L*. The plot of SA increases very gradually for increasing lengths compared to the other models. The linear increase in width with length (proportionate growth) of **model II**, with width *w*=*L*/ *A*_*R*_ suggests that the scaling of area scales quadratically with length *SA* ∝ *L*^2^(from Equation 6). This increase exceeds that of experimental data. If cell width however saturates with increasing length as predicted by **model III**, the surface area is expected to show biphasic behavior- for the initial phase when width increases linearly with the length, *SA* ∝ *L*^2^and when the width reaches maximum it becomes*SA* ∝ *L* (similar to **model I**). The experimental measure of SA with length data matches the predictions of **model III**, with model parameters estimated from data (**Figure 5B**). At the same time, the relative flux rate of a cell through the cell membrane alone is determined by the SA/V ratio. A comparison between model predictions for increasing cell lengths (**Figure 6C**) demonstrates that the relative flux will only change slightly if the cell width is constant (**model I**), rapidly decline if the aspect ratio is constant, (**model II**) and remains constant after an initial decline if the width saturates with length (**model III**). This suggests that maintaining constant cell width could be the optimal strategy for maximizing flux when faced with cell filamentation, i.e. increasing cell length, as compared to a strategy of proportional growth, i.e. linear increase in width with length (**Figure 6D**). At the same time the initial increase in cell width that rapidly saturates, could be the consequence of the molecular network that determines cell size regulation in rod-shaped cells such as *E. coli*.

**Figure 6:**
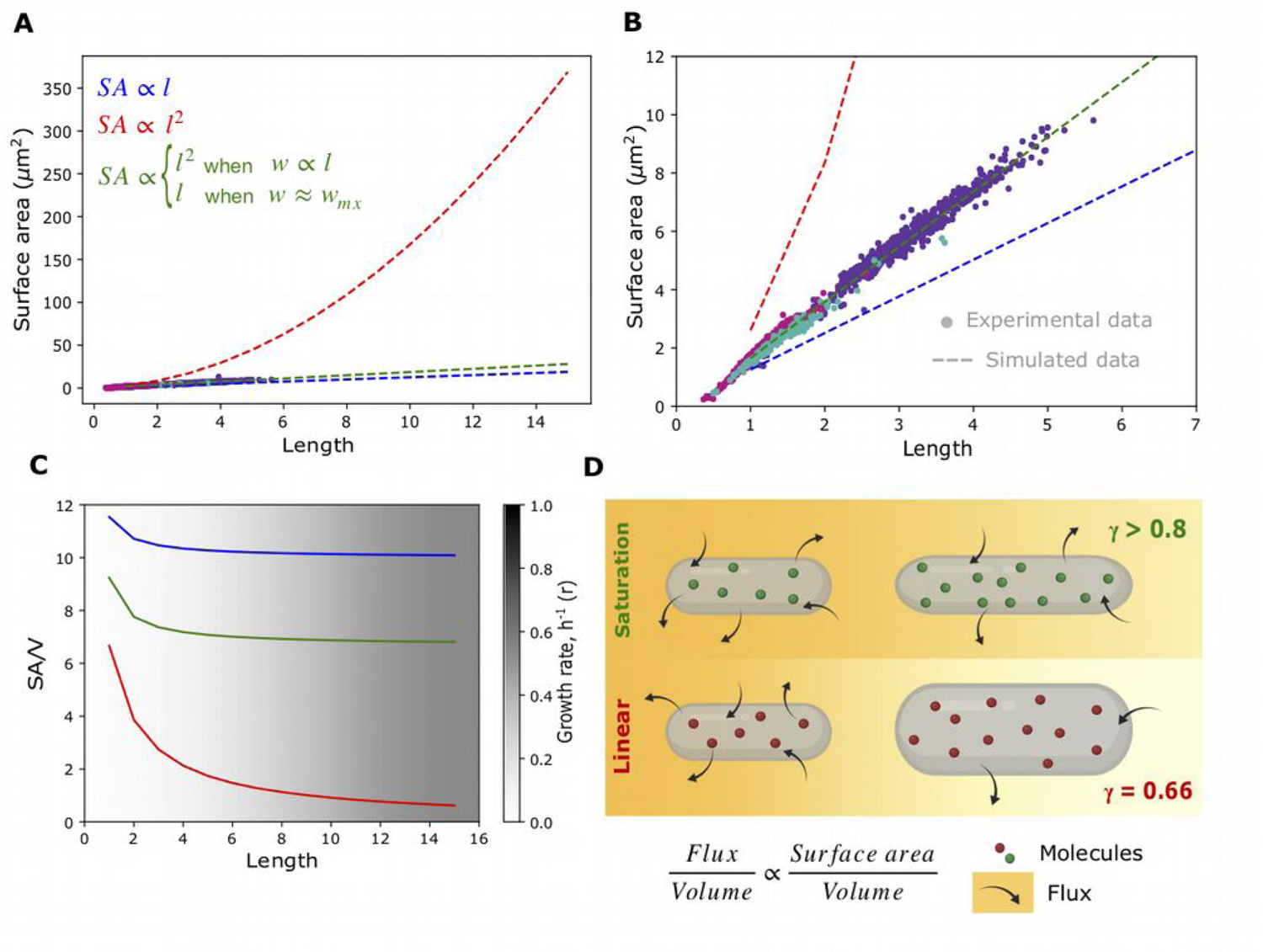
Effect of cell width regulation on surface area and flux for elongated rod-shaped cells. **(A, B)** The surface area (SA) is predicted to scale with cell length for the three alternative models as: (I) *SA* ∝ *L*^2^ (blue), (II) *SA* ∝ *L*^2^(red), (III) *w* ∝ *L*when *SA* ∝ *L*and *SA* ∝ *L* when w = w_mx_ (green) over a large range of cell lengths. The experimental data (circles) is plotted for comparison and **(B)** magnified for 1 to 5 μm cell lengths. Colored circles indicate: *E. coli* MG1655 untreated (magenta) and treated with 100 μg/ml cephalexin (violet) and *E. coli* DH5⍺ (cyan). **(C)** The predicted SA/V are plotted for cell lengths ranging from 1 to 15 μm, with widths determined by models I, II or III. The gradient depicts the increasing growth rates that correspond to the lengths, from Equation 2 (Results section). **(D)** The schematic represents the relative flux for cells with increasing cell length with widths determined by the linear, II (red) or saturation, III (green) models. The SA/V scaling exponent γ is related to the proportionality with flux per unit volume. Green and red circles: molecules, arrows: flux magnitude, orange gradient: magnitude of flux. Schematic created with BioRender.com.

## Discussion

The scaling of surface area and volume is fundamental to comparative physiology and can offer insights into regulatory mechanisms. In this study we find the *E. coli* surface area scales with volume with an exponent of γ ~ 0.8 to 0.9, contrary to reports in the literature that had claimed that such scaling exponents are specific only to certain bacteria and not *E. coli*. This scaling exponent is consistently higher than 2/3 predicted for proportional increase of width and length, irrespective of growth rate, mutations that determine nucleoid dynamics and culture substrate. Statistical averaging of experimental cell volume and area distributions can reproduce the proportional scaling of SA-V reported for *E. coli*, while statistical sampling of length and width distributions confirms SA-V scales with exponents greater than 2/3. The simulation also predicts γ increases from 0.6 to 0.9 with increasing mean cell size, suggesting a role for cell size regulation in scaling behavior. We test three alternative non-molecular models of *E. coli* width regulation with length - constant, proportional and saturation and find experimental data is best fit best by the width saturation model. It also predicts the increase in surface area with length for short cells to be quadratic, while filamentous cells are expected to show a linear increase. This biphasic dependence could have implications for flux maximization, within the constraints of the molecular mechanisms that regulate cell length and width.

Amongst the phenomenological models for cell division, the most relevant to *E. coli* is the adder model (Amir, 2014; Campos et al., 2014). Conventionally, the adder model proposed that a constant length is added to the birth length before the cell can divide, indicating a constant increment to cell length per cell cycle. Microfluidics based cell length analysis has proven to be a powerful tool to examine the dynamics of cell lengths as seen in case of the ‘mother-machine’ (Wang et al., 2010a) that later was used to demonstrate constant cell length increment confirmed the ‘adder’ model (Taheri-Araghi et al., 2015). Recent work demonstrating the limits of measurement has suggested measurement errors and cell size variability could make both the adder model and sizer, i.e. the cell divides when it reaches a specific size, equally likely (Facchetti et al., 2019). Added volume of a cell could arise from an increase in the cell length, width or both. Our model predicts that the increment in volume for short cells will be in terms of both length and width whereas for longer cells it is expected to be only in terms of length. Such a prediction could be tested in future using single cell growth dynamics in microfluidics.

Bacterial cell size was first demonstrated to exponentially increase with growth rate by Schaechter et al. (Koch and Schaechter, 1962; Schaechter et al., 1962). This Schaechter’s nutrient growth law has recently been modified to account for a uniform initiation mass per replication origin independent of growth condition (Si et al., 2017). At a single cell level, the rates of volume growth relative to surface area has been shown to determine cell size based on a predicted homeostasis of SA/V (Harris and Theriot, 2016). This relative growth rate model implies that the surface area growth rate will change depending on a threshold volume growth rate- below the threshold the SA rate will increase and above it decrease (Harris and Theriot, 2018). We believe this is consistent with our findings of a biphasic increase in surface area with length (L), where surface area increases quadratically with length, SA ∝ L^2^, for short cells while it shows a linear increase for longer cells, SA ∝ L, if we consider increasing the mean cell length L to represent increasing growth rate, based on the nutrient growth law. As expected from geometric considerations the ratio of SA/V decreases with increase in length. However when γ ~ 0.6 (~2/3) this decrease is more rapid as compared to the more gradual decrease predicted for cells with a higher scaling exponent (γ ~ 0.8) (**Figure 6C**). The increase in length corresponds by extension to an increase in growth rate based on Schaechter’s nutrient growth law (**Equation 2**). This suggests that systems with higher scaling exponents are more robust to changes in growth rate. This robustness will imply that elongated cells whose widths saturate rather than linearly scale, would be expected to have a higher flux per unit volume due to the relation 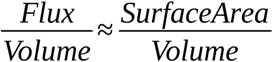 derived from the proportionality of flux with surface area (**Figure 6D**). Therefore we hypothesize that a the observed scaling exponent γ ~ 0.8 seen in *E. coli* could act to minimize the effect of increasing size with growth rate, by regulating cell shape within the limits placed on it by molecular machinery that determines cell shape including membrane synthesis precursors and MreB that is known to regulate cell width (Young, 2010). While these results are obtained with *E. coli* cells, we expect these general principles to also apply to other rod-shaped bacteria.

The actin homolog MreB maintains the rod-shape of most non-spherical bacteria (Bendezú and De Boer, 2008) and mechanically contributes equally to bacterial cell wall stiffness as the peptidoglycan layer (Wang et al, 2010b). The almost linear relation between cell width and MreB inhibitor concentration A22 (Ouzounov et al., 2016) as well as expression level of MreB (Zheng et al., 2016) point to MreB being the primary dose-dependent regulator of *E. coli* cell width. In *Caulobacter* thin cells and wide cells regulate their widths in order to achieve an equilibrium SA/V (Harris et al, 2014). The reduced stiffness of cells with lowered MreB (Zheng et al., 2016) suggests that the shape of the cell is correlated to its ability to withstand pressure. Additionally narrow cells divide in less time with wider cells taking longer to divide, related to FtsZ ring assembly (Liang et al., 2020). This points to multiple roles for MreB as both a tension sensor and a tension creator. However, based on our results the apparent cross-talk between length regulation and width remains to be understood.

In conclusion we find that the shape of *E. coli* cells as described by the surface area scaling with volume for increasing cell size deviates from the ‘ideal’ 2/3 proportional scaling. Simulations and experimental analysis explore the role of two potential mechanisms that could explain it-uncorrelated population variability in cell lengths and widths and regulation of *E. coli* cell width that saturates as a function of cell length. Through a combination of experiments and simulations, we find that the scaling exponent points to a mechanism of cell width regulation for elongated cells. This could have consequences for robust adaptation by *E. coli* to changing growth rates may be relevant for other rod-shaped bacteria.

## Materials and Methods

### Bacterial culture, fixation and staining

Two strains of *E. coli* MG1655 (6300) and DH5, obtained from the Yale CGSC Collection were grown in Luria Bertani (LB) broth (HiMedia, Mumbai, India) at 37°C with constant shaking at 800 rpm (MIR-154-PE, Panasonic Corporation, Japan). Cells were sampled from the mid log phase of growth and stained by addition of FM4-64 (Sigma-Aldrich, Mumbai, India) and grown for 30 min. Cells were washed three times with phosphate buffered saline (PBS) and fixed in 4% (w/v) paraformaldehyde (Sigma-Aldrich, Mumbai, India). The fixed cells were air dried on slides and observed with 60% glycerol mountant. Mid-log phase *E. coli* MG1655 growing in LB were treated with 100 μg/ml Cephalexin (Sigma-Aldrich, Mumbai, India) for 2 hours, and stained and fixed similar to the untreated cells.

### Microscopy and image analysis

Fixed cells of *E.coli* MG1655 and DH5 were acquired using the 100× oil objective (N.A. 0.95) on a Nikon TiE microscope (Nikon Corp., Japan) in fluorescence (Excitation Filter EX40/25, Barrier Filter BA605/55) and DIC channels. and images were obtained. Images were preprocessed using a in-house developed MATLAB (Mathworks Inc., Natick, MA, USA) code, followed by analysis for length and width calculation. Images of FM4-64 stained and fixed cells were adjusted for brightness using *imadjust* (saturated intensity values to top and bottom 1%) and cell outlines were segmented by sequentially detecting edges using the Canny method (Canny, 1986) with threshold 0.2. Gaps were filled and regions bridged to arrive at cell outlines. Cells were filtered based on a minimum solidity criterion between 0.3 and 1 to remove small and irregular objects, with cell boundaries were optimized using an edge-based active contour method (Kass et al., 1988), with a smoothing factor 1 and a contraction bias of −0.1. Optimized contour boundaries were overlaid on the original input image and interactively selected to avoid cell debris and out of focus cells. Selected cells with contours were oriented vertically along their major axis and the midline calculated by iteratively finding the center of mass along the vertical axis. Cell widths were taken to be the means of the 3rd and 4th quartiles of the minimal distance between the midlines and the contour boundaries. By taking only top 50% values we exclude narrow widths at the ends of the cell from analysis.

### Data analysis

Based on cell length and width data, geometric considerations were used to calculate cell surface area (SA) by:

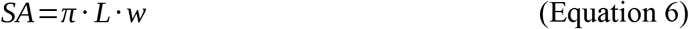

and volume (V) by:

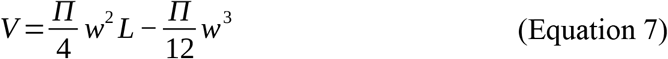

This is based on a spherocylinder assumption with L being the pole-to-pole length and *w* the cell width, similar to previous work (Ojkic et al., 2019). The correlation between surface area (SA) and volume (V) was obtained by fitting the data with Equation 1, with γ the scaling exponent and pre-factor being the free parameters. The aspect ratio (A_R_) was calculated as A_R_=L/w for each data point. In-house developed Python scripts were used to perform the calculations (Python3) using Pandas, NumPy, SciPy packages and Matplotlib (v3.4.2), with polyfit used for curve fitting. Cell SA and V from previously measured *E. coli* MG1655 cell lengths with varied growth rates (Gangan and Athale, 2017; http://dx.doi.org/10.5061/dryad.2bs69) combined with width (w) data that was extrapolated from growth rate (r) correlation *w*=0.14 × 2^0.33·*r*^, as previously described (Taheri-Araghi et al., 2015).

### Sampling cell sizes in a statistical model

Cell lengths were sampled from either lognormal or normal distributions while widths were assumed to be normally distributed, using Numpy (Python3). The input to the distributions involved providing a range of mean values and fixed standard deviations. Mean length values were taken over a range that is observed experimentally (1.1 to 12 μm) the corresponding width (*w*) was calculated from the aspect ratio (A_R_) based on 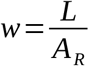 where *L* is the mean length. The aspect ratio was assumed to be constant based on growth-rate dependent mean cell size measurements by Taheri-Araghi et al. (2015). Standard deviations of lengths were calculated from a constant CV (s.d./mean) of 5 (20% of mean length) while width s.d. was 0.035 μm, based on averaging the experimentally reported s.d. of *E. coli* widths. All codes are available on GitHub at https://github.com/CyCelsLab/bacterialsizeScaling.

## Acknowledgements

TK is funded by a PhD fellowship from IISER Pune. DK is supported by a fellowship from the Department of Biotechnology, Govt of India (DBT/JRF/BET-18/I/2018/AL/188) and CNRS-IISER Pune joint PhD program. Neha Khetan is acknowledged for initially pointing out inconsistencies in the surface area and volume scaling in literature. We thank Nishad Matange and Sunish Radhakrishnan for the kind gift of bacterial strains.

## Supplementary material/Supporting Information

### 1. Supporting Results

#### 1.1 Allometric scaling is independent of growth medium and cell stage synchrony

The statistics of cell surface area and volume scaling seen in fixed cell analysis (Methods) could arise from differences in cell cycle stage of the cells, resulting in our observation of the scaling exponent γ > 2/3. We test this hypothesis by using length obtained from microcolonies grown on agar pads (Figure 1A) in order to sample all cell stages. We obtained comparable results using these cell lengths and obtain scaling exponent γ ~ 0. 88 (Figure S1B). Thus the SA-V scaling exponent is consistently greater than 2/3 and independent of cell stage and growth condition.

#### 1.2 *E. coli* mutants for SOS response pathway proteins show comparable SA-V scaling

Cell lengths of three strains of *E. coli* mutant for one of the SOS response and nucleoid occlusion pathway genes- recA, sulA and slmA- were plotted based on previous data (Figure S2A), and widths inferred from growth rates (Equation 2, main text). The SA scales with V by an exponent γ ranging between 0.8 to 0.91 (Figure S2B). The SA and V scaled by their minimal values can be compared for all three strains, and resulted in a scaling exponent of 0.91 (Figure S2C).

### 2. Supporting Figures

**Figure S1.**
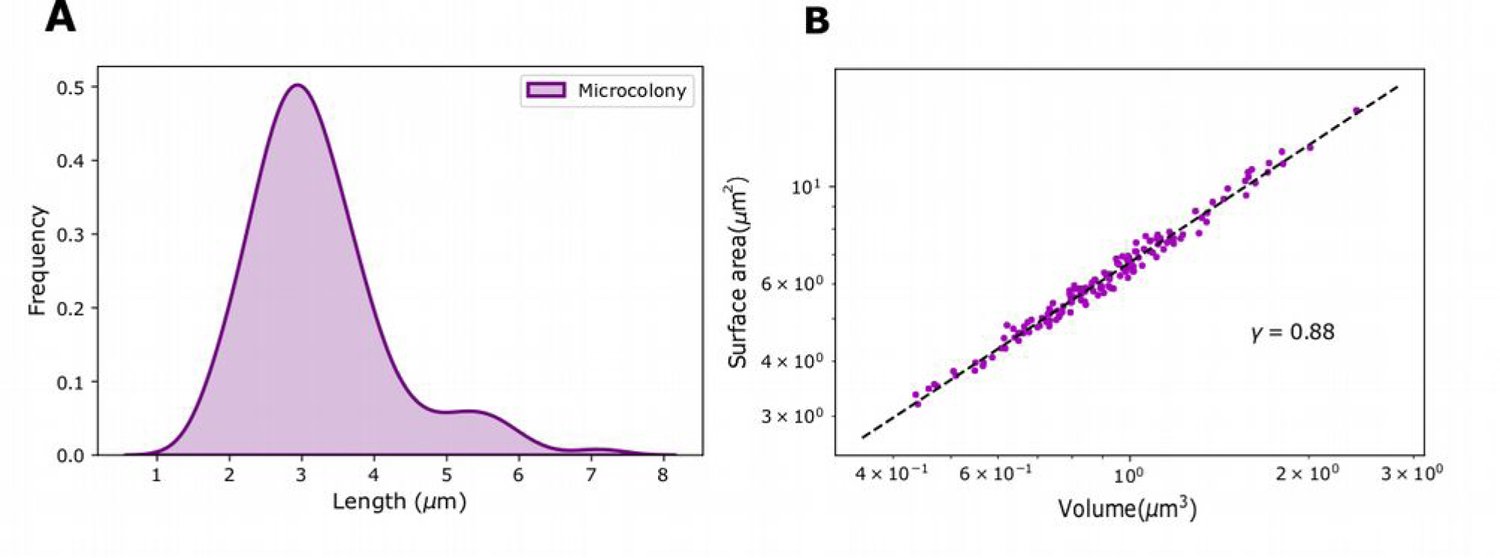
SA-V scaling of *E. coli* grown on agar-pads. **(A)** Cell lengths from microcolonies grown on LB agar-pads were used to estimate **(B)** surface area and volume (circles). Data was fit to Equation 1 to estimate γ. Data is taken from a previous report (Gangan and Athale, 2017) and widths were measured as described in the Materials and Methods section.

**Figure S2:**
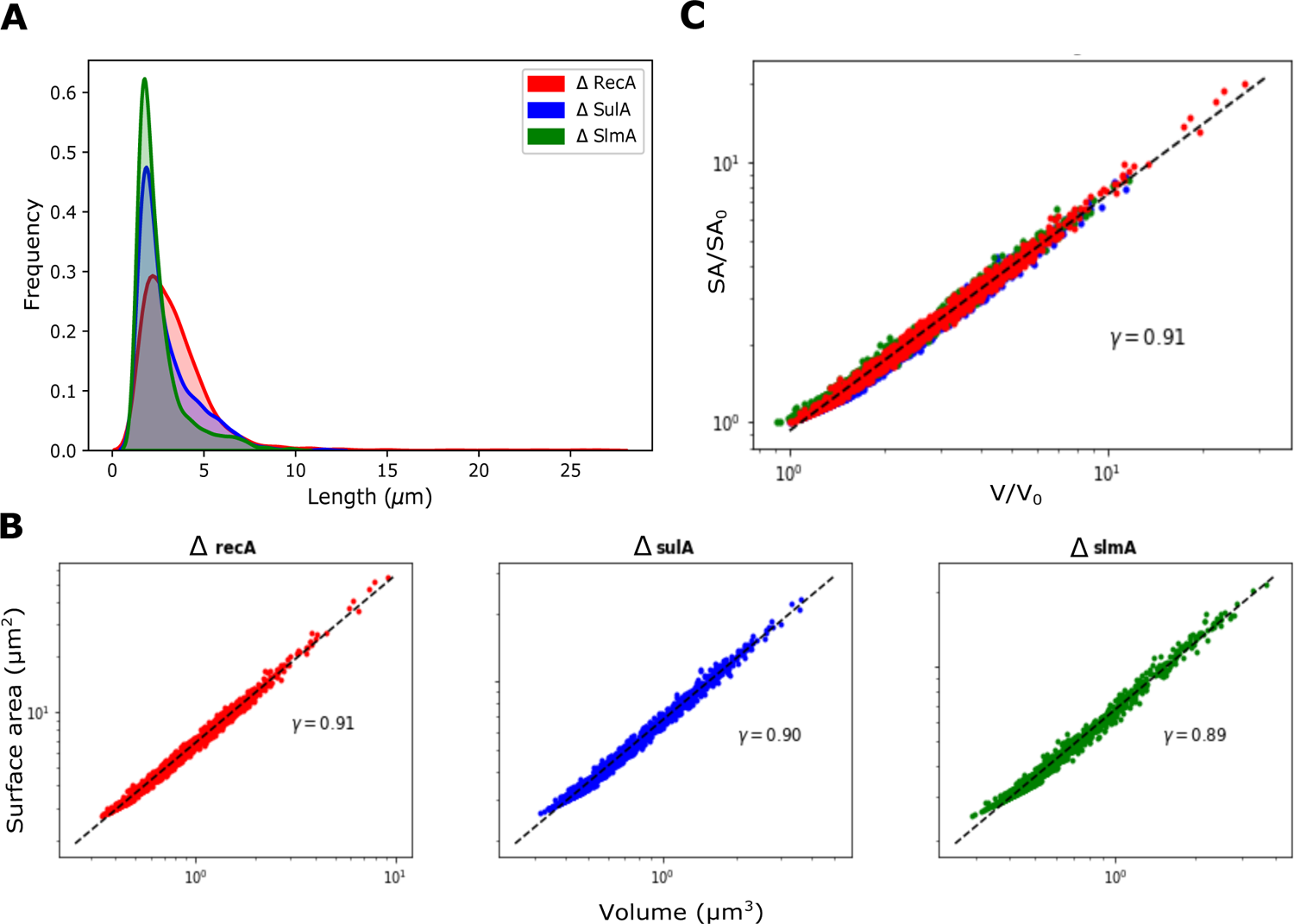
Effect of mutants associated with the nucleoid on *E. coli* scaling. **(A)** Cell length frequency histograms of *E. coli* mutants ∆recA (red), ∆sulA (blue), ∆slmA (green) were used to **(B)** correlate the surface area (SA) and volume (V) and fit to Equation 2 to obtain the scaling exponent γ. **(C)** The SA and V data for the mutants was normalized by the corresponding smallest values (S_0_, V_0_) and the scaling exponent was inferred similarly. All the data is taken from previous reported cell lengths measured from DIC images (Gangan and Athale, 2017).

**Figure S3.**
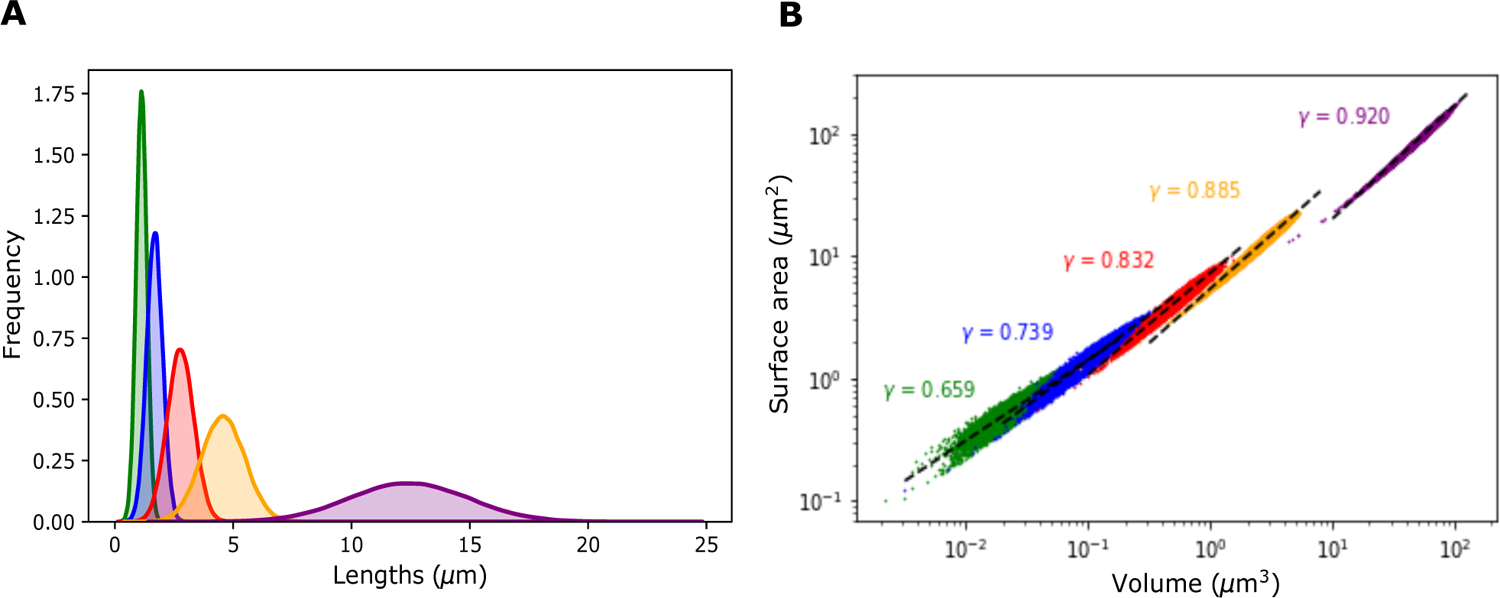
Simulating the SA-V scaling of normally distributed cell lengths. **(A)** Simulated cell length distributions that follow a normal distribution for increasing mean lengths (μ_L_) and standard deviation (_L_) values were combined with normally distributed widths (Figure 3C, main text) to generate (B) a plot of surface area with volume and fit to Equation 1 to obtain γ. Colors indicate increasing mean length values.

